# Machine-Learning-Based spike marking in signal and source space EEG from a patient with focal epilepsy

**DOI:** 10.64898/2026.03.06.710063

**Authors:** Lala Jafarova, Demet Yeşilbaş, Christoph Kellinghaus, Gabriel Möddel, Stjepana Kovac, Stefan Rampp, Daniela Czernochowski, Sebastian Sager, Ayşegül Güven, Turgay Batbat, Carsten H. Wolters

**Affiliations:** Institute for and Biosignalanalysis, University of Münster; Department of Cognitive Science, RPTU Kaiserslautern-Landau; Department of Biomedical Engineering, Graduate School of Natural and Applied Sciences, Erciyes University, Kayseri, 38039 Turkey; Epilepsy Center Münster-Osnabrück, Department of Neurology, Klinikum Osnabrück, Osnabrück, Germany; Epilepsy Center Münster-Osnabrück, Department of Neurology with Institute of Translational Neurology, University Hospital Münster, Münster, Germany; Department of Neurosurgery, University Hospital Erlangen, Erlangen, Germany; Department of Biomedical Engineering, Faculty of Engineering, Erciyes University, Kayseri, 38030 Turkey; Department of Mathematics, Otto von Guericke University, Magdeburg, Germany

**Author notes:** These authors contributed equally to this work (Lala Jafarova, Demet Yeşilbaş).

**Keywords:** interictal epileptiform discharges, EEG, Artificial Neural Networks, Source Localization, Epilepsy

## Abstract

Accurate detection of interictal epileptiform discharges (IEDs) in electroencephalography (EEG) plays a crucial role in epilepsy diagnosis. Our work investigates the classification of IEDs using Artificial Neural Networks (ANNs) trained on EEG data represented in both signal and source space. Source waveforms were computed using equivalent current dipole models fitted using either a 1-parameter fixed-orientation or a 3-parameter projection approach, both localized to a single best-fit position during the rising flank of the IED. The ANN was trained on raw and feature-extracted versions of signal space and source space data. Feature extraction significantly improved performance across all domains. The highest accuracy (0.98) was achieved in signal space using Katz Fractional Dimension (KFD). In source space analyses, the 1-parameter and 3-parameter models achieved a maximum accuracy of 0.84, with statistical features performing best for the fixed-orientation model and KFD for the free orientation model. Additionally, annotations from three independent expert markers showed considerable variability, with ANN performance falling within the range of inter-expert agreement. These findings support the potential of ANN-based tools to assist expert evaluation in future clinical workflows.

## 1. Introduction

Epilepsy is a chronic neurological disorder that affects more than 50 million people worldwide. It is characterized by a persistent tendency toward unprovoked, recurrent seizures, caused by abnormal or excessive neuronal activity that leads to changes in behavior, sensation, or motor function (Fisher et al., 2014). With appropriate diagnosis and treatment, seizure freedom can be achieved in approximately 70% of individuals; however, epilepsy can also impact cognitive, psychological and social functioning (Milligan, 2021; Sarmast et al., 2020).

Clinical diagnosis begins with the classification of seizures as either focal (originating in one brain area) or generalized (affecting both hemispheres simultaneously), which is essential for selecting appropriate treatment (Chavan & Desai, 2023; Hussein et al., 2018; Sarmast et al., 2020). Standard diagnostic approaches involve clinical evaluation, electroencephalography (EEG) to identify abnormal brain activity and often neuroimaging (MRI) to detect structural abnormalities (Milligan, 2021). In clinical practice, epilepsy patients who are candidates for surgical intervention often undergo prolonged EEG monitoring. Accurate identification of epileptic spikes is critical for localizing the epileptogenic zone and guiding surgical decisions, as misclassifications may directly affect treatment outcomes. Even in single-patient scenarios, clinical decisions rely heavily on EEG findings, making reliable automated spike detection essential.

EEG data analysis can be conducted in two complementary domains. Signal-space analysis examines the potential differences measured by scalp electrodes that arise from electrical activity in the brain. While straightforward, this approach is affected by volume conduction, which mixes signals and can complicate the interpretation of underlying neural sources. In contrast, source-space analysis applies forward and inverse modeling techniques to estimate the cortical generators of the measured EEG signals. By incorporating biophysical head model and source constraints, such as approaches, can provide improved spatial interpretability of neural activity (Kirsch et al., 2006; van de Steen et al., 2016). Source-space representations have been shown to be particularly useful for the analysis of transient events, such as IEDs (Barzegaran & Knyazeva, 2017).

EEG is a fundamental, non-invasive neurophysiological tool for diagnosis and classification. It records the electrical potentials generated by synchronized neural populations in the cerebral cortex via electrodes placed according to a standardized system (Mecarelli, 2019). Volume conduction enables these cortical sources to be measurable at the scalp; however, accurate interpretation depends on appropriate modeling of head tissue conductivity and source configurations. EEG can capture both ictal (occurring during seizure) and interictal (occurring between seizures) activity, making it indispensable for distinguishing epileptic from non-epileptic events (Rosenow & Lüders, 2001). Interictal epileptiform discharges (IEDs), such as spikes and sharp waves, are used to localize the irritative zone, whereas ictal patterns help identify the seizure onset zone. Generalized seizures often exhibit bilaterally synchronous spike-and-wave discharges, while focal seizures are typically associated with localized interictal activity (Noachtar & Rémi, 2009).

Despite its widespread clinical utility, conventional scalp EEG has limited spatial resolution due to the smoothing effects of volume conduction and is highly sensitive to accurate modeling of head tissue conductivities in source reconstruction (Gross et al., 2021; Vorwerk et al., 2024). Source localization approaches, including dipole fitting methods, aim to estimate the underlying neural generators from scalp recordings. Such techniques improve spatial interpretability by incorporating forward modeling of head conductivity and source configurations(Gross et al., 2021). These methodological developments provide a foundation for more reliable quantitative analyses of epileptiform activity and motivate the use of advanced computational approaches for automated EEG assessment.

IEDs are paroxysmal transient abnormalities, such as sharp, high-amplitude peaks, that occur in the EEG between seizures (de Moraes & Callegari, 2016). Accurate detection of IEDs is crucial for diagnosing epilepsy and avoiding misdiagnosis, which occurs in 20% to 30% of referred patients (Benbadis, 2009; Ramakrishnan et al., 2025). According to the International Federation of Societies for Electroencephalography and Clinical Neurophysiology (IFSECN), these epileptiform patterns are transients distinguishable from background activity, with a spiky morphology and are important for localizing and complexity: a spike lasts 20-70 ms, a sharp wave lasts 70-200 ms and they can combine with slow waves to form complexes (Noachtar et al., 1999).

Morphological analysis of IEDs can be quantified using scoring systems, such as the Bergen Epileptiform Morphology Score (BEMS), which weights features like spike slope and amplitude to improve diagnostic accuracy (Aanestad et al., 2023). However, spike detection is challenging due to the low signal-to-noise ratio, variable morphology and subjectivity in visual annotation. This has led to the development of numerous automated algorithms that use signal processing and pattern recognition to enhance consistency (Holleman et al., 2011; Kim & Kim, 2000).

More recently, sophisticated Machine Learning models, such as Artificial Neural Networks (ANNs), which learn directly from EEG signals to improve the accuracy and generalizability of IED detection (Kumar & Upadhyay, 2025; Ma et al., 2024). In addition to traditional amplitude- and morphology-based measures, recent studies suggest that epileptiform spikes exhibit complex, nonlinear and transient dynamics. Consequently, nonlinear and complexity-based descriptors such as entropy, fractal dimensions and spectral features have been shown to provide discriminative information for automated spike detection (Bagheri et al., 2019). These features capture complementary aspects of spike morphology that may not be apparent in simple amplitude or duration measurements, potentially improving classification performance when combined with machine learning approaches.

Building on these differences, the present study investigates whether EEG features derived from source-space representations provide complementary or superior information to traditional signal-space features for automated interictal spike detection. We hypothesize that training artificial neural networks on source-space EEG data improves the reliability of spike detection compared to signal-space analysis alone. To test this hypothesis, EEG signals are filtered and annotated, MRI-based head models and source localization is performed to obtain source waveforms. ANN-based classifiers were then trained and evaluated using features extracted separately from signal-space and source-space representations.

A key distinguishing aspect of this study is the availability of annotations from three experienced epileptologists who independently reviewed extended EEG recordings. This enables not only the development of automated detection methods but also the comparison of inter-rater variability with the performance of ANN-based approaches.

## 2. Materials and Methods

### 2.1. Ethics Statement

The study was conducted according to the guidelines of the Declaration of Helsinki and approved by the institution’s ethical review boards (Ethik Kommission der Ärztekammer Westfalen-Lippe und der Westfälischen Wilhelms-Universität Münster, 25.05.2021, Ref. No. 2021-290-f-S). The patient gave written informed consent for her data to be used in anonymized form for scientific publications.

### 2.2. Data Collection

EEG data were collected from a 26-year-old female patient with refractory focal epilepsy originating in the left frontal lobe near Broca’s area, treated at Münster University Hospital between 2018 and 2023. The patient experienced non-motor seizures characterized by impaired thinking and communication, occurring from weekly up to four times daily. Despite multiple anti-seizure medications, the condition remained pharmacoresistant, prompting a presurgical evaluation in 2018 (Antonakakis et al., 2024). As the initial invasive EEG did not localize the seizure onset zone, a second invasive evaluation using depth electrodes (iEEG) was performed in autumn 2021. Subsequently, craniotomy and resection under awake conditions were conducted in January 2023. Additional clinical details can be found in Section S2.2 and S2.10 of the Supplementary Material in Antonakakis et al. (2024).

In total, 22 hours of scalp EEG were recorded using a 19-channel 10-20 system at a sampling rate of 200 Hz and segmented into 22 one-hour sessions. All analyses were performed in MATLAB R2025b using FieldTrip^1^ (Oostenveld et al., 2011), the Signal Processing Toolbox^2^, the Statistics and Machine Learning Toolbox^3^and some custom scripts.

### 2.3. EEG preprocessing

EEG preprocessing was performed to ensure high signal quality for reliable spike detection. Recordings from 19 clinical channels of the international 10-20 system were analyzed, with particular relevance to the left frontal region given the F3-centered IEDs investigated in this study (Hinrichs et al., 2020). For visual annotation by epileptologists, the EEG were reviewed using bipolar montages over the left frontal region to facilitate identification of phase reversals associated with focal activity. For computational analysis, continuous EEG recordings were bandpass filtered between 0.5 and 40 Hz to suppress slow drifts and high-frequency noise. The cleaned recordings, totaling 22 hours, were segmented into non-overlapping 1-second epochs (200 samples at a sampling rate of 200 Hz), yielding 79,876 epochs for subsequent analysis.

Interictal epileptiform discharges were manually annotated in the preprocessed EEG by three independent epileptologists (EP1, EP2 and EP3). Each reviewer independently labeled epochs they judged to contain IEDs, resulting in 2,890 spike epochs identified by EP1, 1.646 by EP2 and 10,279 by EP3. This variability reflects the well-documented subjectivity of visual EEG interpretation. All remaining epochs were labeled as non-spike, leading to pronounced class imbalance with non-spike epochs substantially outnumbering spike epochs.

To examine the impact of labeling strategy and reduce subjectivity, two complementary label sets were constructed. An AND set included only epochs labeled as spikes by all three epileptologists, providing high-certainty spike annotations. An OR set included epochs marked by at least one reviewer, increasing sensitivity to subtle epileptiform activity at the cost of greater label uncertainty. These labeling strategies formed the basis for subsequent signal-space analysis, feature extraction and ANN model training.

### 2.4. Signal Space Representation

In the signal-space representation, EEG data were analyzed directly at the scalp electrode level using the preprocessed recording described above. Each non-overlapping 1-second epoch (200 samples at 200Hz) from the 19 electrodes of the international 10-20 system was treated as an independent observation. No spatial transformation or source modeling was applied in this representation and the recorded potentials reflect the mixed contributions of underlying cortical generators as measured at the scalp.

Signal-space epochs labeled according to the AND and OR annotation strategies served as the baseline data for feature extraction and ANN-based spike classification. This representation constitutes the conventional approach used in clinical EEG analysis and provides the reference against which the source-space representation was compared in terms of spike detection performance.

### 2.5. Source Localization

Source localization was performed by transforming scalp EEG into a source-level representation using a realistically shaped three-compartment (scalp, skull, brain) head volume conductor model for the EEG forward problem. Spike epochs identified using the AND and OR labeling strategies were averaged separately and temporally aligned to the negative peak at electrode F3, where maximal spike amplitude was observed, in order to obtain representative spike waveforms for source analysis. Non-spike epochs were retained as unaligned 1-second segments.

Grand-average waveforms were then inspected to identify a stable interval for dipole fitting (Figure 1). The early rising phase (*−*20 to *−*5 ms relative to the F3 peak) was selected, as it more closely reflects the primary epileptic generator and is less influenced by propagation effects that may dominate at the spike peak (Antonakakis et al., 2024; Aydin et al., 2017, 2015; Lantz et al., 2003; Mălîia et al., 2016; Plummer et al., 2019).

**Figure 1.**
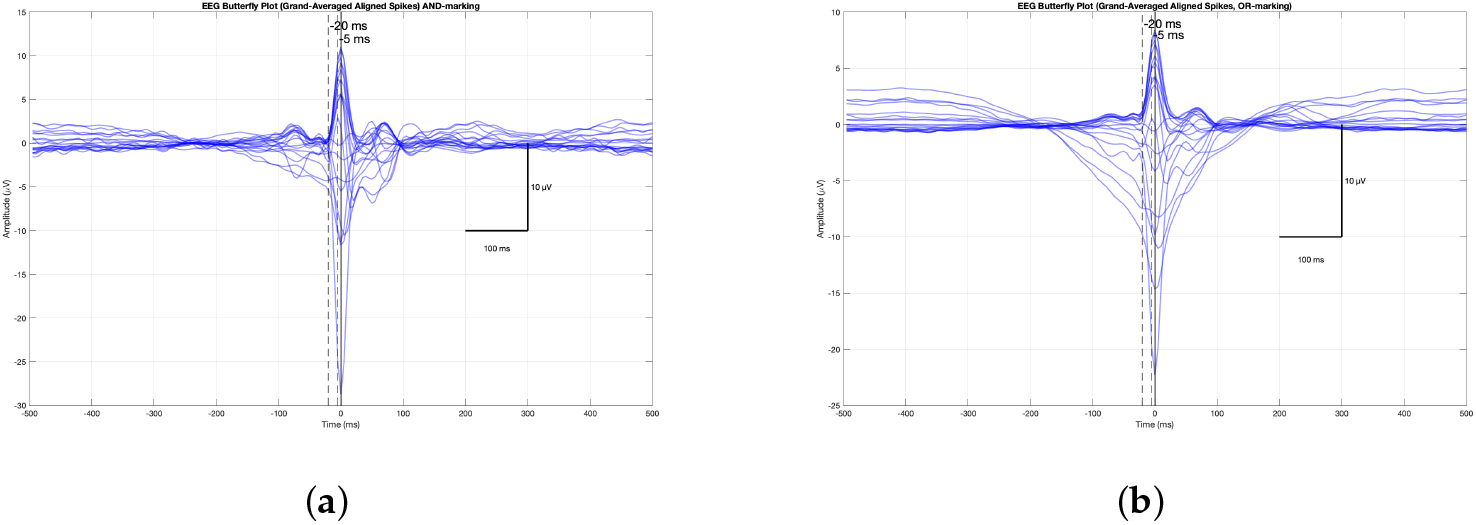
Grand average of IED epochs for (**a**) conservative AND-marking (unanimous expert agreement) and (**b**) sensitive OR-marking (at least one expert) strategies. Each colored trace represents one of 19 EEG channels in average reference. Waveforms are temporally aligned to the negative peak amplitude at the F3 electrode (time = 0 ms, vertical solid line). The vertical dashed lines at *−*20 ms and *−*5 ms indicate the time window used for subsequent dipole fitting and source reconstruction. Time axis: *−*100 ms to +100 ms relative to peak. Amplitude axis: *µ*V.

An individual T1-weighted MRI was used to construct a realistically shaped three-compartment (scalp, skull, brain) head volume conductor model for symmetric BEM EEG forward modeling implemented via OpenMEEG in FieldTrip (Gramfort et al., 2010). Standard 10-20 electrode positions were aligned to the scalp surface and a 3-mm source grid was defined within the brain volume. The resulting lead field matrix provided the forward mapping between cortical sources and scalp potentials. All steps were visually inspected to ensure anatomical accuracy and spatial consistency prior to inverse modeling (Brette & Destexhe, 2012).

A single moving equivalent dipole (ECD) was fitted to the grand-average spike waveforms for both labeling strategies within the selected time window. Both AND- and OR-labeled averages yielded high goodness-of-fit values, indicating that the modeled dipoles accounted for most of the observed variance. Dipoles were localized consistently to the left frontal lobe near F3, in agreement with the clinically identified seizure onset zone (Antonakakis et al., 2024).

Based on the dipole location, source nodes were selected in two stages: (1) an anatomically constrained left frontal region for fixed-orientation (1D) projection and (2) a localized patch of ten nodes surrounding the dipole for evaluating the effect of orientation parameterization. For each of these nodes, source waveforms were computed using three orthogonal orientations (3D, yielding 3 × 10030) (Figure 2), we computed the L2-norm (vector magnitude) of the three-component waveform at each node: 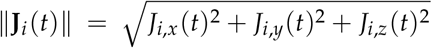. Source waveforms were obtained by solving a least-squares inverse problem,

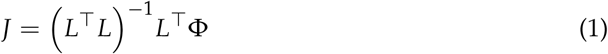

where *λ* is the Tikhonov regularization parameter, selected automatically using the L-curve criterion to balance residual error and solution smoothness for patch-level projection.

**Figure 2.**
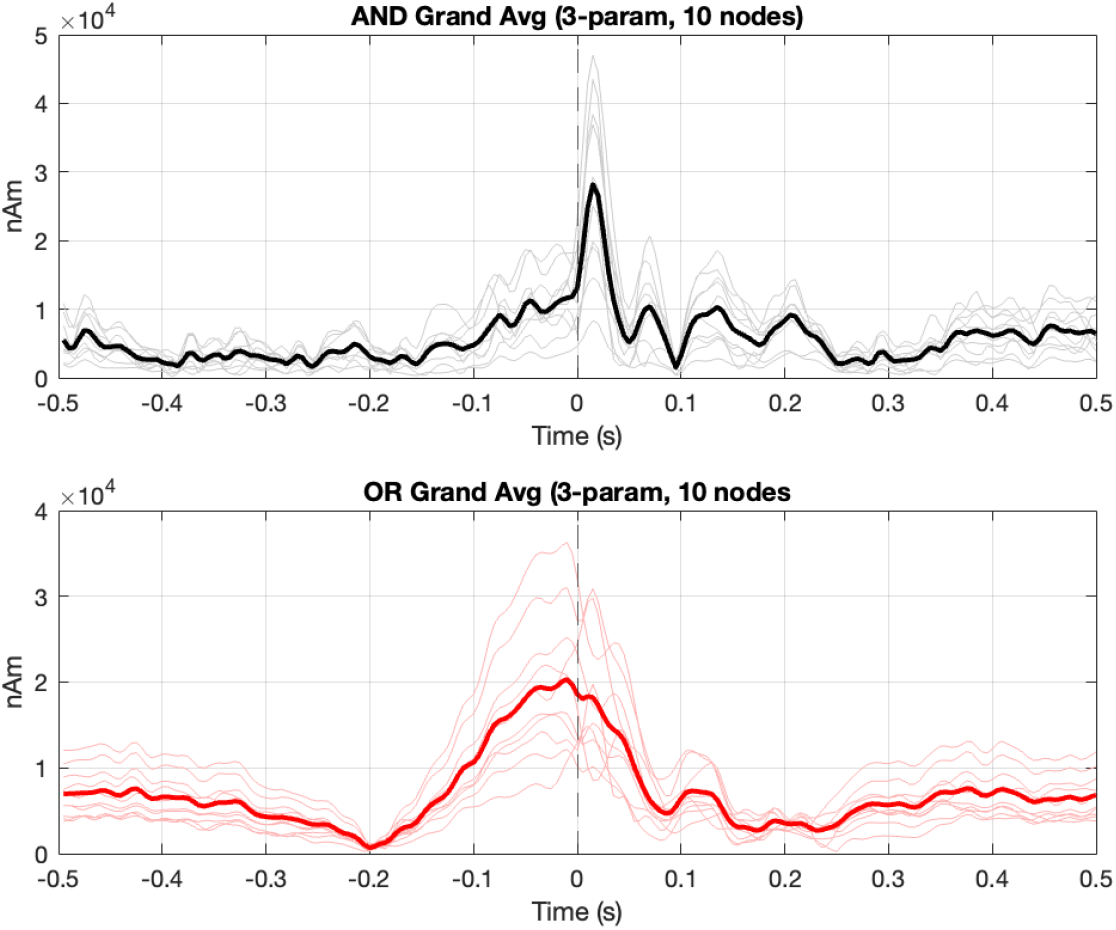
Patch vel source waveforms for grand-average IEDs. The plots show the vector magnitude 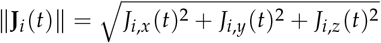 of the three-component (*x, y, z*) source estimates at each node, expressed in nanoampere-meters (nAm), averaged across the 10 closest cortical nodes to the fitted dipole location for (**a**) AND-marked and (**b**) OR-marked d ata. Gray l ines: individual node magnitudes; black line (a) / red line (b): mean across nodes.

### 2.6. Dataset Balancing

Following signal space representation and source space reconstruction, class imbalance between spike and non-spike epochs was addressed prior to feature extraction. Severe class imbalance can bias machine learning models toward the majority class and distort subsequent feature estimation, particularly for rare events such as interictal epileptiform discharges (Ruiz de Miras, 2016).

To mitigate this effect, spike and non-spike epochs were randomly down-sampled to create balanced datasets with equal class representation. This procedure was applied separately for AND and OR labeling strategies. The resulting balanced epoch sets were then used as input for subsequent feature extraction in both signal and source space and for ANN model training. This approach ensured unbiased feature distributions and improved model generalization (Chawla et al., 2002; Güven et al., 2020).

### 2.7. Feature extractions

To prepare data for ANN-based epileptic spike detection, feature extraction was applied to transform 19-channel EEG recordings from both signal and source spaces representation into informative, low dimensional feature vectors. EEG data were segmented into non-overlapping 1-second epochs (19 channels × 200 samples), a duration chosen to capture the full temporal characteristics of interictal epileptiform discharges, using labels provided by three independent epileptologists.

A set of well-established statistical, nonlinear, and frequency-domain measures commonly used to characterize EEG signal dynamics (Yeşilbaş et al., 2025). Statistical features included the mean, standard deviation, skewness, and kurtosis of the amplitude distribution. Nonlinear complexity features comprised fractal dimensions (KFD and Higuchi), the Lyapunov exponent quantifying dynamical instability and entropy-based measures (approximate entropy and Shannon entropy) assessing signal unpredictability and irregularity. Frequency-domain features were computed as band power within conventional EEG sub-bands: delta (0.5-4 Hz), theta (4-8 Hz), alpha (8-13 Hz), beta (12-30 Hz) and gamma (30-60 Hz) (Altınkaynak et al., 2024)

The same feature extraction procedure was applied to both signal space and source space data to enable a direct and fair comparison of their respective contributions to automated interictal spike detection.

### 2.8. Artificial Neural Networks

Artificial Neural Networks (ANNs) are computational models inspired by the biological nervous system, where simplified artificial neurons compute weighted sums of inputs, apply a nonlinear activation function and generate outputs (Guilhoto, 2018). A feed-forward ANN with one hidden layer of 10 neurons and a single output neuron was used in this study for binary spike classification, chosen for its ability to model nonlinear patterns in moderately sized EEG datasets.

To separate feed-forward ANN models with identical architectures were trained using MATLAB’s *patternnet*^4^. By default, training was performed using the Scaled Conjugate Gradient (SCG) backpropagation algorithm, which efficiently incorporates second-order curvature information without explicit Hessian computation or line search. All inputs were z-score normalized prior to training to ensure comparable scaling across features.

**Signal-space model:** The raw input representation used 3800-dimensional vectors (19 channels × 200 time points) from average-referenced EEG epochs. **Source-space model:** For the 1-parameter projection, the input consisted of 2000-dimensional vectors (10 nodes × 200 time points). For the 3-parameter projection, the input consisted of 6000-dimensional vectors (30 components × 200 time points), preserving the three orthogonal current directions at each node.

Model performance was evaluated through 10-fold cross-validation to ensure robust generalization and prevent overfitting (Chavakula et al., 2013; Paul & Karn, 2023). Within each fold, the training data were further subdivided (70% training, 30% testing) for internal model selection. Classification performance was evaluated using multiple complementary metrics, including accuracy, sensitivity, specificity, precision, F1-score, Geometric mean (G-mean),and Cohen’s Kappa. Accuracy measures overall correctness, while sensitivity and specificity reflect the correct identification of spikes and non-spike predictions and F1-score balances precision and recall (Boonyakitanont et al., 2019; Guglielmi et al., 2025). The G-mean assesses the balance between sensitivity and specificity, making it useful for imbalanced datasets. Cohen’s Kappa evaluates agreement beyond chance, with values above 0.6 indicating substantial agreement (Nandy et al., 2019; Sabharwal, 2021)

## 3. Results

This section presents the classification performance of the ANN under different data representations and preprocessing strategies, with the primary goal of evaluating the impact of feature extraction and source space transformation on IED detection.

ANN performance was assessed for four conditions: raw signal-space EEG, feature-extracted signal-space EEG, raw source-space representation and feature-extracted source-space data. Results are reported separately for AND- and OR-marking strategies to account for differences in annotation certainty.

### 3.1. Signal-space classification

Using raw signal-space EEG epochs without feature extraction resulted in near-chance classification performance for both marking strategies (Table 1). Accuracies were 0.52 (AND) and 0.51 (OR), barely exceeding random guessing. Cohen’s kappa values were low, indicating agreement only slightly better than chance. Sabharwal (2021). Sensitivity and specificity were unbalanced, with neither metric consistently exceeding 0.60, demonstrating that raw average-referenced EEG waveforms alone are insufficient for reliable ANN-based IED detection.

**Table 1.**
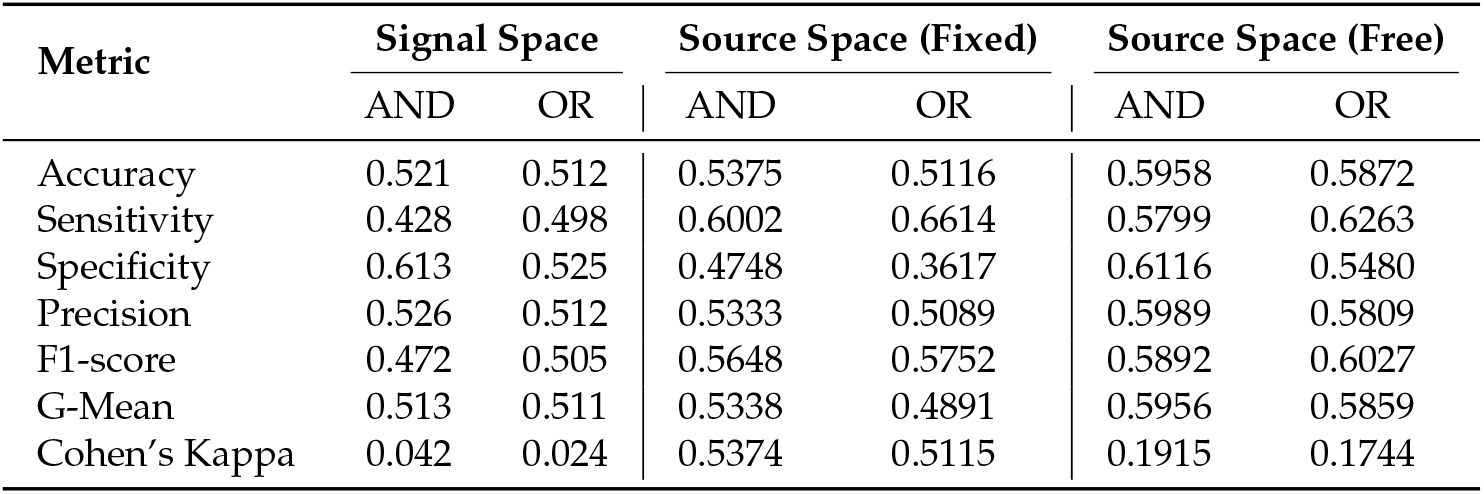
ANN classification performance for raw EEG and source-space data across AND- and OR-marking strategies.

In contrast, feature extraction led to a dramatic improvement in classification performance (Table 2). For AND-marked data, several feature sets achieved accuracies above 0.97, with balanced sensitivity and specificity exceeding 0.95, indicating balanced performance across both spike and non-spike classes. Most notably, the Katz Fractal Dimension (KFD, F2) alone reached accuracy of 0.98 (Cohen’s *κ* = 0.96, indicating almost perfect agreement), comparable to more complex combined feature sets such as F6 (Statistical + KFD + Power Spectral Density (PSD): 0.98), F8 (KFD + Lyapunov: 0.98) and F11 (KFD + Approximate Entropy: 0.97).

**Table 2.**
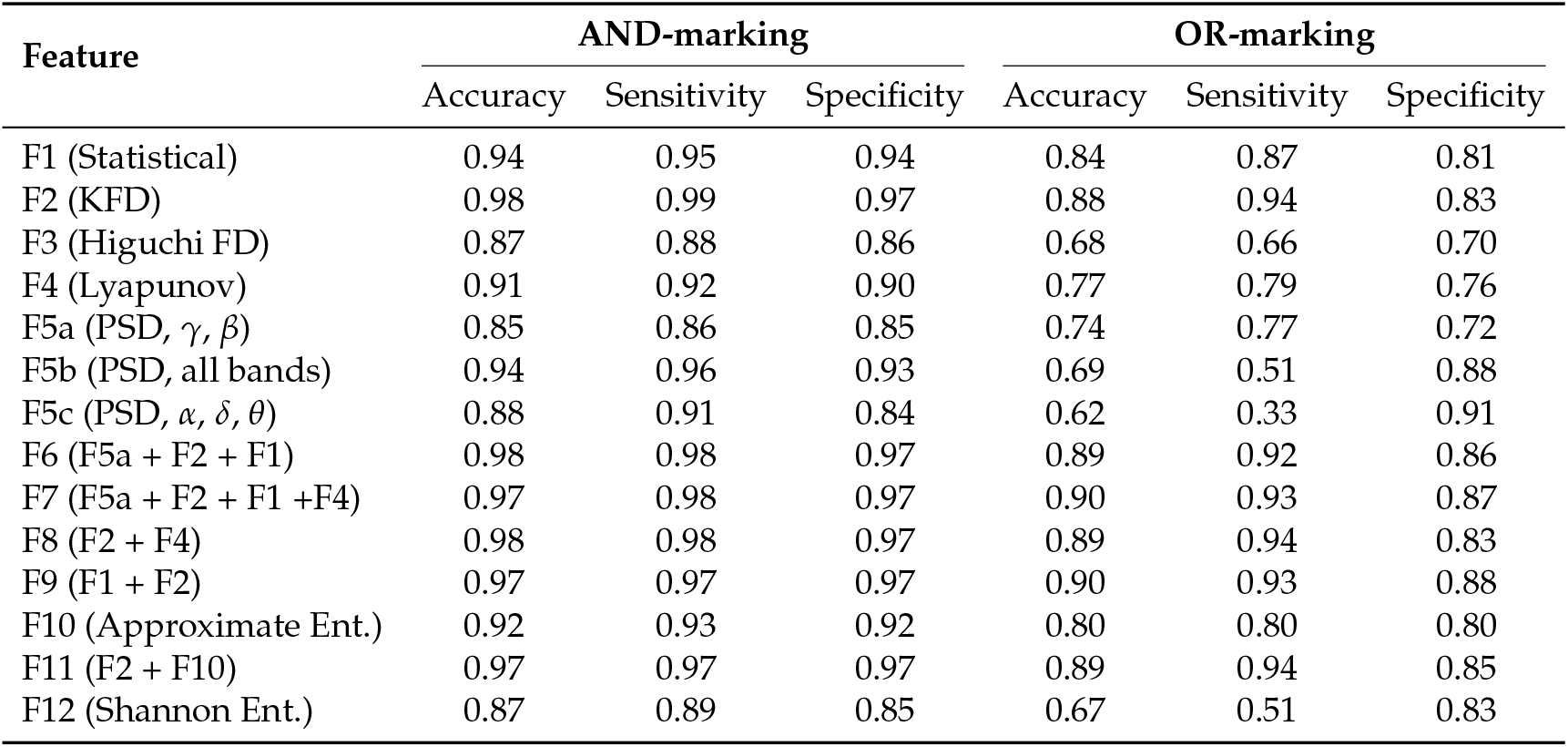
Signal Space ANN Classification Performance for AND- and OR-marking.

| Feature | AND-marking |  |  | OR-marking |  |  |
| --- | --- | --- | --- | --- | --- | --- |
|  | Accuracy | Sensitivity | Specificity | Accuracy | Sensitivity | Specificity |
| F1 (Statistical) | 0.94 | 0.95 | 0.94 | 0.84 | 0.87 | 0.81 |
| F2 (KFD) | 0.98 | 0.99 | 0.97 | 0.88 | 0.94 | 0.83 |
| F3 (Higuchi FD) | 0.87 | 0.88 | 0.86 | 0.68 | 0.66 | 0.70 |
| F4 (Lyapunov) | 0.91 | 0.92 | 0.90 | 0.77 | 0.79 | 0.76 |
| F5a (PSD, $\gamma$, $\beta$) | 0.85 | 0.86 | 0.85 | 0.74 | 0.77 | 0.72 |
| F5b (PSD, all bands) | 0.94 | 0.96 | 0.93 | 0.69 | 0.51 | 0.88 |
| F5c (PSD, $\alpha$, $\delta$, $\theta$) | 0.88 | 0.91 | 0.84 | 0.62 | 0.33 | 0.91 |
| F6 (F5a + F2 + F1) | 0.98 | 0.98 | 0.97 | 0.89 | 0.92 | 0.86 |
| F7 (F5a + F2 + F1 + F4) | 0.97 | 0.98 | 0.97 | 0.90 | 0.93 | 0.87 |
| F8 (F2 + F4) | 0.98 | 0.98 | 0.97 | 0.89 | 0.94 | 0.83 |
| F9 (F1 + F2) | 0.97 | 0.97 | 0.97 | 0.90 | 0.93 | 0.88 |
| F10 (Approximate Ent.) | 0.92 | 0.93 | 0.92 | 0.80 | 0.80 | 0.80 |
| F11 (F2 + F10) | 0.97 | 0.97 | 0.97 | 0.89 | 0.94 | 0.85 |
| F12 (Shannon Ent.) | 0.87 | 0.89 | 0.85 | 0.67 | 0.51 | 0.83 |

Performance for OR-marked data was lower overall (best accuracy: 0.90) but still substantially improved compared to raw signal space (0.51). Best-performing feature sets achieved accuracies between 0.88 and 0.90, with F2 again leading at 0.88 accuracy, closely followed by F6, F7 and F9 (Statistical + KFD). Cohen’s kappa values ranged from 0.76 to 0.80, indicating substantial agreement. The systematically lower performance for OR-marking reflects the increased morphology variability and subjectivity inherent in this labeling scheme, where spikes are marked by at least one epileptologist.

These results demonstrate that appropriate feature engineering transforms near-random performance (0.51-0.52) into clinically meaningful classification (0.88-0.98). Notably, in both results and those reported by (Yeşilbaş et al., 2023), KFD alone achieved performance levels close to the best feature combinations, suggesting that KFD is the primary driver of high accuracy, while combinations offer only marginal additional benefit.

### 3.2. Source-space classification

Applying ANN classification directly to source-localized waveforms without feature extraction yielded only modest improvements over raw signal space (Table 1). The fixed-orientation model showed an achieved accuracy of 0.53 (AND) and 0.51 (OR), barely above chance. The free-orientation model achieved slightly higher accuracies of 0.59 (AND) and 0.58 (OR). However, performance remained substantially lower than that achieved using feature-extracted signal-space data, indicating that the dimensionality reduction by source analysis alone does not improve the classification much.

Feature extraction in source space improved classification performance relative to raw source data (Tables 3 and 4), but peak accuracies remained lower than those obtained in signal space. For the fixed-orientation model with AND-marked data, the maximum accuracy of 0.84 was achieved with statistical features (F1) and mixed feature sets performing best (F6, F7). For the free-orientation (3-parameter) model, best performance increased to 0.85 (F6, F7) and F2 being 0.84, representing only a modest improvement despite preserving full three-dimensional source orientation information.

**Table 3.** Source Space ANN Classification Performance for Fixed-Orientation Model (AND- and OR-marking).

| Feature | AND-marking |  |  | OR-marking |  |  |
| --- | --- | --- | --- | --- | --- | --- |
|  | Accuracy | Sensitivity | Specificity | Accuracy | Sensitivity | Specificity |
| F1 (Statistical) | 0.84 | 0.81 | 0.86 | 0.65 | 0.55 | 0.74 |
| F2 (KFD) | 0.68 | 0.62 | 0.73 | 0.59 | 0.59 | 0.60 |
| F3 (Higuchi FD) | 0.62 | 0.72 | 0.51 | 0.59 | 0.67 | 0.52 |
| F4 (Lyapunov) | 0.59 | 0.65 | 0.52 | 0.57 | 0.60 | 0.53 |
| F5a (PSD, $\gamma$, $\beta$) | 0.77 | 0.67 | 0.86 | 0.50 | 0.37 | 0.63 |
| F5b (PSD, all bands) | 0.78 | 0.72 | 0.85 | 0.50 | 0.65 | 0.36 |
| F5c (PSD, $\alpha$, $\delta$, $\theta$) | 0.59 | 0.75 | 0.43 | 0.51 | 0.64 | 0.37 |
| F6 (F5a + F2 + F1) | 0.84 | 0.81 | 0.87 | 0.64 | 0.59 | 0.70 |
| F7 (F5a + F2 + F1 + F4) | 0.84 | 0.83 | 0.85 | 0.65 | 0.59 | 0.71 |
| F8 (F2 + F4) | 0.66 | 0.66 | 0.66 | 0.58 | 0.58 | 0.59 |
| F9 (F1 + F2) | 0.83 | 0.82 | 0.85 | 0.65 | 0.57 | 0.73 |
| F10 (Approximate Ent.) | 0.61 | 0.63 | 0.59 | 0.56 | 0.62 | 0.50 |
| F11 (F2 + F10) | 0.69 | 0.69 | 0.68 | 0.58 | 0.61 | 0.55 |
| F12 (Shannon Ent.) | 0.63 | 0.80 | 0.45 | 0.53 | 0.88 | 0.16 |

**Table 4.** Source Space ANN Classification Performance for Free-Orientation Model (AND- and OR-marking).

| Feature | AND-marking |  |  | OR-marking |  |  |
| --- | --- | --- | --- | --- | --- | --- |
|  | Accuracy | Sensitivity | Specificity | Accuracy | Sensitivity | Specificity |
| F1 (Statistical) | 0.75 | 0.73 | 0.77 | 0.64 | 0.59 | 0.69 |
| F2 (KFD) | 0.84 | 0.84 | 0.84 | 0.68 | 0.59 | 0.69 |
| F3 (Higuchi FD) | 0.63 | 0.68 | 0.58 | 0.61 | 0.67 | 0.56 |
| F4 (Lyapunov) | 0.63 | 0.70 | 0.57 | 0.60 | 0.63 | 0.57 |
| F5a (PSD, $\gamma$, $\beta$) | 0.80 | 0.75 | 0.85 | 0.54 | 0.80 | 0.28 |
| F5b (PSD, all bands) | 0.82 | 0.76 | 0.87 | 0.54 | 0.84 | 0.25 |
| F5c (PSD, $\alpha$, $\delta$, $\theta$) | 0.53 | 0.90 | 0.25 | 0.53 | 0.90 | 0.16 |
| F6 (F5a + F2 + F1) | 0.85 | 0.84 | 0.86 | 0.65 | 0.60 | 0.69 |
| F7 (F5a + F2 + F1 + F4) | 0.85 | 0.84 | 0.86 | 0.67 | 0.64 | 0.70 |
| F8 (F2 + F4) | 0.82 | 0.81 | 0.82 | 0.63 | 0.65 | 0.61 |
| F9 (F1 + F2) | 0.73 | 0.71 | 0.75 | 0.60 | 0.58 | 0.63 |
| F10 (Approximate Ent.) | 0.65 | 0.68 | 0.63 | 0.60 | 0.62 | 0.58 |
| F11 (F2 + F10) | 0.83 | 0.72 | 0.83 | 0.64 | 0.66 | 0.61 |
| F12 (Shannon Ent.) | 0.68 | 0.78 | 0.58 | 0.53 | 0.83 | 0.22 |

Notably, KFD (F2), which achieved 0.98 accuracy i signal space, showed substantially reduced performance in source space: 0.68(1D) and 0.84 (3D) for AND marking. This suggests that the source projection, while improving spatial specificity, may compress spatiotemporal signal complexity critical for certain nonlinear features. The free-orientation model partially recovered performance (0.84 vs 0.68), supporting the importance of preserving orientation dynamic for complexity-based features, though the improvement remained modest compared to the signal-space benchmark.

For OR-marked data, overall performance was further reduced, with best accuracies around 0.65 (fixed) and 0.68 (free), compared to 0.90 in signal space. This decline highlights the sensitivity of source-space feature-based classification to labeling uncertainty and increased variability.

Overall, feature extraction was the dominant factor in improving classification in both signal and source spaces (from 0.52 to 0.84-0.98), but signal-space features outperformed source-space features. While source localization offers improved spatial interpretability, it did not enhance classification performance. The free-orientation model provided only marginal gains over fixed orientation (0.85 vs 0.84 for AND-marking).

## 4. Discussion

This is the first study where 22 hours of continuous EEG data, independently marked by three experienced epileptologists, were used to systematically train and evaluate artificial neural networks with and without feature extraction in signal space and source space. By comparing raw waveform classification against feature-based approaches across multiple representation domains (signal space, 1-parameter source projection and 3-parameter source projection), we provide a comprehensive assessment of the methodological factors that determine automated spike detection performance. Our results demonstrate that feature extraction is essential for reliable ANN-based IED detection, while raw EEG data in both signal and source space lacks sufficient discriminative structure for accurate classification without dimensional reduction through carefully designed features.

In signal space, the application of morphological, spectral and nonlinear features significantly improved performance, achieving accuracies up to 0.98 for AND-marked data and 0.90 for OR-marked data. Raw signal-space waveforms, by contrast, achieved only chance-level performance (accuracy ≈ 0.52), demonstrating that high-dimensional EEG time series require feature engineering to expose spike-specific patterns to conventional classifiers.

Katz Fractal Dimension (KFD) emerged as the most discriminative single feature, achieving 0.98 accuracy (AND marking) and 0.88 accuracy (OR-marking) without combination with other features. This finding is consistent with the pioneering work of Yeşilbaş et al. (2023), who demonstrated that KFD achieves near optimal classification performance (0.96 AND-marking, 0.87 OR-marking) when applied to the 12 most informative bipolar channels selected from a standard 19-electrode montage. Our results extend these findings by showing that KFD maintains its discriminative power even when applied to average-referenced data across all 19 channels, achieving comparable performance, suggesting that the choice of referencing scheme has minimal impact on KDF’s discriminative ability once appropriate features are extracted.

The strong performance of fractal dimension measures for spike detection aligns with broader literature on nonlinear EEG analysis. Arle and Simon (1990) showed that fractal dimension could distinguish transient epileptiform events from background EEG based on self similar temporal structure. Accardo et al. (1997) showed that Higuchi and Katz fractal dimensions effectively quantify the complexity changes associated with pathological brain states. Our finding that KFD substantially outperforms Higuchi FD (0.98 vs 0.87 accuracy for AND marking) is consistent with Anisheh and Hasanpour (2009), who noted that KFD requires less computational complexity while maintaining discriminative power for abrupt waveform changes characteristic of epileptic spikes.

Statistical features (mean, standard deviation, skewness, kurtosis) also achieved strong performance (0.94 accuracy for AND-marking), consistent with Shoka et al. (2019), who identified these features among the optimal extraction methods for epilepsy classification across multiple studies. The combination of KFD with statistical features (F9) achieved 0.97 accuracy, demonstrating complementarity between fractal and moment-based descriptors-fractal dimension captures self-similar temporal structure, while statistical moments quantify amplitude distribution properties.

Spectral features based on conventional EEG sub-bands (delta, theta, alpha, beta, gamma) showed variable performance depending on band selection. Beta and gamma bands (F5a: 0.85 accuracy) outperformed all five bands combined (F5b: 0.94), suggesting that higher-frequency components contain spike-specific information, consistent with the known spectral signature of sharp transients. These findings align with Hassan et al. (2019), who reported that higher-frequency sub-bands yielded better epileptic seizure detection performance using multi-band feature extraction with feed-forward neural networks.

Entropy-based measures (approximate entropy: 0.92, Shannon entropy: 0.87) showed moderate discriminative power, though they underperformed relative to fractal dimension. This contrasts with some prior work Swami et al. (2016) that reported excellent seizure detection using entropy features combined with general regression neural networks. The discrepancy may reflect differences between interictal spike detection and ictal seizure detection, as seizures involve more pronounced state changes that entropy measures are designed to quantify.

The comparison between AND-marking and OR-marking strategies revealed that data consistency plays a critical role in model performance, which may reflect the well-documented variability and subjectivity of visual EEG interpretation reported in literature (Barkmeier et al., 2012). AND-marked data, representing events unanimously identified by all three epileptologists, yielded substantially higher classification accuracies (up to 0.98) compared to OR-marked data (up to 0.90). OR-marked data, which includes events identified by at least one expert, introduced greater variability due to its inclusion of ambiguous and heterogeneous spike morphologies. Nevertheless, even under this noisier labeling condition, feature-based representations remained superior to raw input, emphasizing the robustness of feature extraction.

Source-space analysis provided insight into the trade-off between anatomical specificity and signal information preservation. Raw source-projected data showed only modest improvements over raw signal-space data (accuracies 0.54-0.60), suggesting that dimensionality reduction through projection alone is insufficient. When features were applied, source-space performance improved substantially, reaching accuracies up to 0.85 (3D projection, AND-marking). However, source-space results did not surpass those obtained in signal space (0.98), indicating that the simplified source model and spatial regularization inherent in inverse modeling limited the amount of exploitable spatiotemporal information. This phenomenon is expected given the ill-posed nature of the EEG inverse problem, where regularization can reduce spatial and temporal resolution and suppress high-frequency components (Grech et al., 2008).

A key finding was the dependence of feature effectiveness on data representation. While KFD dominated in signal space, statistical features became more effective in the 1-parameter source projection. This shift can be explained by the smoothing and noise-suppressing effect of source localization, which produces simpler, more “peaky” waveforms that are well captured by basic statistical measures but less suitable for complexity-based features consistent with previous reports noting that inverse source reconstruction involves spatial regularization and smoothing, which may attenuate fine temporal details of scalp EEG activity (Michel & Brunet, 2019). The improved performance of KFD under the free-orientation (3-parameter) source model supports this interpretation (F2: 0.84 vs. 0.68 for 1-parameter), as additional degrees of freedom preserve more dynamic information, such as waveform rotation and propagation from sulcal to the gyral regions.

These findings align with previous work showing that feature extraction is fundamental for IED detection and that more complex source models better capture epileptiform dynamics (Kaviri & Vinjamuri, 2024; Singh et al., 2022). Unlike studies using distributed source models, the simplified projections used here restrict the representational power of the source-space. This limitation explains why source location, despite its anatomical advantages, did not outperform signal-space analysis in classification accuracy.

### 4.1. Limitations and Future Directions

The most significant limitation of this study is the single-patient design. While 22 hours of continuous EEG provide substantial within-patient variability, epileptic spikes vary considerably across individuals depending on epilepsy etiology, cortical origin and medication status (Noachtar et al., 1999). Multi-patient validation is essential, as previous studies show performance degradation when moving from single-patient to multi-patient scenarios (Johansen et al., 2016; Tjepkema-Cloostermans et al., 2018). Methodologically, our 3-compartment symmetric boundary element method (symBEM) with moving dipole fits could be improved using calibrated 6-compartment finite element models with anisotropic conductivities (Höltershinken et al., 2025) and distributed inverse methods such as beamforming our minimum-norm estimation (Hämäläinen & Ilmoniemi, 1994), which my preserve more discriminative source space information. The feed-forward ANN architecture was intentionally simple for fair comparisons by modern convolutional neural networks (CNN)(Johansen et al., 2016). Finally, clinical validation studies are essential to evaluate whether automated spike detection reduces epileptologist workload while maintaining diagnostic accuracy in real-world clinical workflow.

## 5. Conclusion

This study investigated ANN-based IED detection using EEG data in both signal and source space, with and without feature extraction. The results indicate that feature extraction is indispensable for accurate classification, whereas raw EEG input alone is insufficient. In signal space, the feature-based model achieved the highest performance, reaching accuracies of up to 0.98, with Katz Fractal Dimension and combined spectral-fractal features proving especially effective.

Although source localization provides valuable anatomical information, its integration with a simplified projection model offered only modest gains and consistently outperformed signal-space performance. The dependence of feature effectiveness on data representation highlights the importance of matching feature design to the underlying signal model. In particular, more complex source models with greater degrees of freedom appear to help preserve the spatiotemporal dynamics of IEDs and fully exploit source-space analysis.

These findings reinforce the critical role of data representation and feature selection in EEG-based IED detection. From a clinical perspective, feature-based automated systems operating in signal space may offer as assistive tools for improving efficiency and consistency in epilepsy diagnostics. Future work should extend this framework to multi-patient datasets, explore richer and more flexible source localization strategies, and investigate a deep learning architecture capable of directly learning spatiotemporal patterns from both signal and source space data.

## Author Contributions

**Lala Jafarova**: Methodology, Software, Formal analysis, Investigation, Data curation, Writing – original draft, Writing – review & editing, Visualization. **Demet Yeşilbaş**: Methodology, Software, Investigation, Writing – review & editing. **Christoph Kellinghaus**: Resources, Data curation (marking of epileptic activity), Writing – review & editing, Supervision. **Gabriel Möddel**: Resources, Data curation (marking of epileptic activity), Writing – review & editing. **Stjepana Kovac**: Resources, Data curation (marking of epileptic activity), Writing – review & editing. **Stefan Rampp**: Resources, Data curation (marking of epileptic activity), Methodology, Writing – review & editing. **Daniela Czernochowski**: Supervision, Writing – review & editing, Funding acquisition. **Sebastian Sager**: Writing – review editing. **Ayşegül Güven**: Supervision, Writing – review & editing, Funding acquisition. **Turgay Batbat**: Methodology, Software, Writing – review & editing. **Carsten H. Wolters**: Conceptualization, Methodology, Resources, Writing – review & editing, Supervision, Project administration, Funding acquisition.

All authors have read and agreed to the published version of the manuscript.

## Institutional Review Board Statement

The study was conducted in accordance with the Declaration of Helsinki and approved by the Institutional Ethics Review Board (Ethik-Kommission der Ärztekammer Westfalen-Lippe und der Westfälischen Wilhelms-Universität Münster, approval date: 25 May 2021, Ref. No. 2021-290-f-S).

## Informed Consent Statement

Written informed consent was obtained from the patient involved in the study.

## Data Availability Statement

All analysis code and processing pipelines are publicly available on GitHub at https://github.com/cafaroval/ML_spike_detection and all data is available upon request.

## Use of Artificial Intelligence

During the preparation of this work, Grammarly for grammar and style checking and Google NotebookLM for literature organization were used.

## Acknowledgments

The authors would like to thank Andreas Wollbrink for his essential technical support. We also thank Yvonne Buschermöhle and Malte Höltershinken from the SIM-NEURO work-group (Institute for Biomagnetism and Biosignalanalysis, University of Münster) for their constructive discussions and support.

## Disclaimer/Publisher’s Note

The statements, opinions and data contained in all publications are solely those of the individual author(s) and contributor(s) and not of MDPI and/or the editor(s). MDPI and/or the editor(s) disclaim responsibility for any injury to people or property resulting from any ideas, methods, instructions or products referred to in the content.

## Footnotes

1 https://www.fieldtriptoolbox.org

2 https://www.mathworks.com/products/signal.html

3 https://www.mathworks.com/products/statistics.html

4 https://www.mathworks.com/help/deeplearning/ref/patternnet.html

